# A novel lethal cuticular structural protein, AaCPR100A and its upstream interaction protein, G12-like, function in cuticle and egg shell formation in the yellow fever mosquito, *Aedes aegypti*

**DOI:** 10.1101/2023.02.05.527159

**Authors:** Jing Chen, Yuchen Wu, Haoran Lu, Gong Cheng, Zhijian Jake Tu, Chenghong Liao, Qian Han

## Abstract

AaCPR100A is a structural protein, found in the soft cuticle of *Ae. Aegypti*. RNAi of *AaCPR100A* resulted in high mortality in *Ae. Aegypti* and abnormal egg development in the surviving mosquitoes. Over thirty proteins that could interact with AaCPR100A were screened out by yeast two-hybrid assay, and subsequently, further verification by hybrid and GST pull-down assays identified that G12-like had the strongest interaction with AaCPR100A. RNAi of *G12-like* suggested it may be related to larva development. Interestingly, the adults in which the *G12-like* gene *was* knocked down were sensitive to low temperature, and their egg shell formation, production, and hatching were affected. *G12-like* has the opposite effect in the upstream expression of *AaCPR100A*, promoting *AaCPR100A* function in the larval stage and inhibiting *AaCPR100A* in the adult stage. In all, functional studies of AaCPR100A and its interaction protein G12-like provide insight into its involvement in cuticle development and formation and egg shell formation.

## Introduction

The insect cuticle is indispensable to insect life *(Polerstock et al., 2002*; *Willis and Muthukrishnan, 2010)*. Its functions include resisting external environmental damage, forming, and maintaining body shape, and enabling motion and life activity *(Gallot et al., 2010*; *Jasrapuria et al., 2012*; *Zhou et al., 2017*). *Ae. aegypti* undergoes complete metamorphosis in its life cycle, and cuticle formation, differentiation, digestion and reconstruction are parts of this process *(Charles, 2010*; *Srivastava et al., 2014*; *Vargas et al., 2014*; *Xie et al., 2022*; *Zhang et al., 2020)*. Cuticular proteins (CPs) affect the structure and mechanical properties of the cuticle *(Charles, 2010*; *Srivastava et al., 2014*; *Vargas et al., 2014*; *Xie et al., 2022*; *Zhang et al., 2020)*. Some structural CPs can induce apoptosis in upregulated genes, causing structural abnormalities in epidermal cells and endocuticle *(Xiong et al., 2017*; *Zhao et al., 2019)*. The largest CP family in arthropods is the CPR family *(Asano et al., 2019*; *Qiao et al., 2020)*. RR-2 proteins are involved in the construction of the outer cuticle (hard cuticle) and RR-1 proteins are involved in the formation of the endocuticle (flexible cuticle) *(Cornman and Willis, 2008*; *Shahin et al., 2016, 2018*; *Vannini and Willis, 2017*; *Zhang et al., 2020)*.

RNAi of CYP4g1 (aldehyde oxidative decarbonylase P450) and CPR in *Drosophila melanogaster* has been found to block the biosynthesis of hydrocarbons in crimson cells *(Qiu et al., 2012)*. Silencing a cuticular gene that contains the RR-1 motif in locusts causes thickening of the inner cuticular layer of larvae. Silencing CPR that contains the RR-1 motif in silkworms reduces the expansion and elasticity of larvae cuticles, making the larva sbody tight and making it unable to crawl, bend or grip *(Qiao et al., 2014)*. Upregulation of CPR genes in *Cryptopygus antarcticus* and *Onychiurus arcticus* in cold regions showed that CPR genes affect cold stress resistance in insects *(Purac et al., 2008*; *Teets and Denlinger, 2014)*.

The insect cuticle consists primarily of three layers. From outer to inner, these are the epicuticle, the exocuticle, and the endocuticle and epidermal cells. In the formation of the insect cuticle, the epidermal cells secrete the epicuticle, exocuticle and endocuticle to form the mature cuticle *(Locke et al., 2001)*. In most insects, the epicuticle is covered with a lipid layer. Two types of lipids, structural lipids and free lipids, are found in the insect cuticle.

Structural lipids participate in formation of the rigid and hard insect cuticle and their amount and distribution determine the differences in rigidity of the insect cuticle across species and layers *(Wigglesworth, 1970)*. Free lipids, on the other hand, are present in the epicuticle. They form a barrier against desiccation *(Hadley, 1981)* that varies in composition and may contain several chemical classes such as alcohols, aldehydes, esters, fatty acids, glycerides, hydrocarbons and ketones. Studies have shown that fatty acid content affects insect ability to withstand cold environments. Lipids and derivatives in the cuticle retain water and decrease cuticle permeability to prevent the invasion of external harmful substances. Lipids, especially cuticular hydrocarbons (CHC), also act as a semichemical, thus affecting insect communication, defense, mating, reproduction and other behaviors *(Howard and Blomquist, 2005*; *Tillman et al., 1999)*. Wang et al. used eosin, a hydrophilic dye, to study the regional distribution and osmotic characteristics of lipids on the body surface of *Drosophila* and showed that lipid content varied across the cuticle *(Wang et al., 2016)*. Lipids constitute approximately 35% of the dry weight of *Ae. aegypti* eggs, most of which is derived from body fat stores and are transported to the ovaries through the hemolymph (Alabaster et al., 2011).

Ae. aegypti is a widely distributed mosquito species in the family Culicidae. Like other holometabolous insects, the larval stage undergoes several molts before pupating and emerging as an adult *(Brust, 1968*; *Vargas et al., 2014*; *Yu et al., 2021)*. AaCPR100A is a cuticular protein of *Ae. aegypti* that is a member of the CPR family containing the RR-1 motif. It was identified using the *Ae. aegypti* genome and RNA-seq data analysis. This suggests it may mainly function in the soft cuticle region. RNA interference of this gene was lethal, which indicates that the gene is important in the development of *Ae. aegypti*. It is of interest to know how the gene functions and how it affects cuticle formation: whether it affects the formation of chitins or lipids in the cuticle, and whether partner proteins are needed to complete their formation. In this study, we investigated gene expression, protein localization, interaction proteins, and phenotypes of gene silenced mosquitoes using RNAi-mediated gene knockdown to resolve these issues. The results identify a new target for developing an innovative mosquito control technique.

## Results

### Localization and relative quantification of AaCPR100A

A primary antibody generated against AaCPR100A weakly imaged this protein in different tissues in the head and more strongly in the thorax (Figure 1a-d). AaCPR100A protein was also widely present in various parts of the cuticle immediately surrounding various tissues. Red fluorescence was more prominent in the abdominal cuticle and the ovarian intersegmental membrane of female adults (Figure 1e-h).

**Figure 1.**
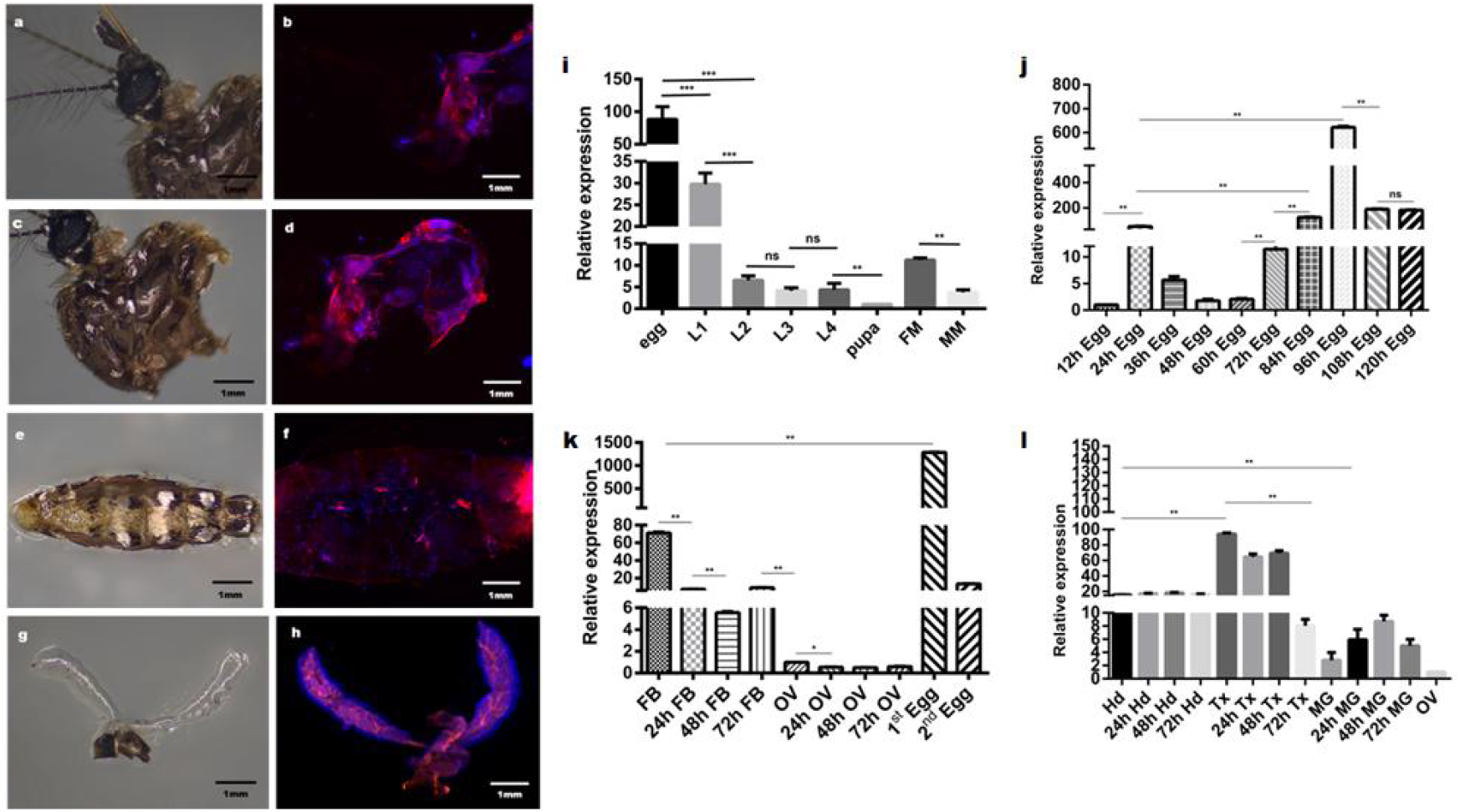
Immunohistochemical analysis of AaCPR100A protein and the transcriptional levels of *AaCPR100A* gene in *Ae. aegypti*. (a) Anatomical head of the mosquito. (b) Localization of AaCPR100A protein in the head. (c) Anatomical thorax of the mosquito. (d) Localization of AaCPR100A protein in the thorax. (e) Anatomical fat body of the mosquito. (f) Localization of AaCPR100A protein in the fat body. (g) Anatomical ovary of the mosquito (h) Localization of AaCPR100A protein in the ovary. (i) Temporal transcription levels of AaCPR100A in different developmental stages. (j) Temporal transcription levels of *AaCPR100A* at different times of egg development; the times 12 h, 24 h and others are times after egg deposition. (k and l) The expression of AaCPR100A in different tissues of adults before and after blood meal; 24 h, 48 h and 72 h are times after blood meal. Abbreviations: L1, first-instar larvae; L2, second-instar larvae; L3, third-instar larvae; L4, fourth-instar larvae; FM, female adults; MM, male adults; FB, fat body; OV, ovary; 1st Egg, eggs that were laid after the first blood meal; 2nd Egg, eggs that were laid after the second blood meal; Hd, head; Tx, thorax; MG, midgut; ns, *P* ≥0.05; *, *P* <0.05; **, *P* <0.01; ***, *P* <0.001.

The transcript abundance of AaCPR100A decreased gradually during development with relative expression highest in the egg stage and the first instar (Figure 1i). Expression was relatively higher in adult females than in adult males (Figure 1i). Transcript abundance of AaCPR100A was greatest in the egg, followed by the thorax and fat body (Figure 1k, l). After blood meal, the AaCPR100A transcription level in the thorax, fat body and ovary was significantly downregulated (Figure 1k, l). The newly laid eggs often hatched in 2-3 d. The newly laid eggs were collected every 12 h, and the greatest transcription abundance of AaCPR100A was observed in the late stage of egg development (Figure 1j).

### RNAi of *AaCPR100A* is lethal and affects egg melanization, delays spawning time and decreases hatching rate

Two methods, immersion and nanoparticle feeding, were used to interfere with the gene in larvae in order to further verify the effect of AaCPR100A RNAi on the larva. The interference of AaCPR100A increased larva lethality compared to the control groups, and lethality gradually increased as the concentration of dsRNA increased (Figure 2a). Figure 2a shows that after 0.01 mg/mL dsRNA immersion interference, the survival rate was 61.7% by 9 d, which was significantly different from the control groups (P < 0.0001). After dipping in 0.1 mg/mL dsRNA, almost all larvae died. There was no significant difference between DEPC water immersion and gus dsRNA treatments with different gus dsRNA concentration (P = 0.63). Chitosan/dsRNA nanoparticles increased the stability of dsRNA (Figure 2b).

**Figure 2.**
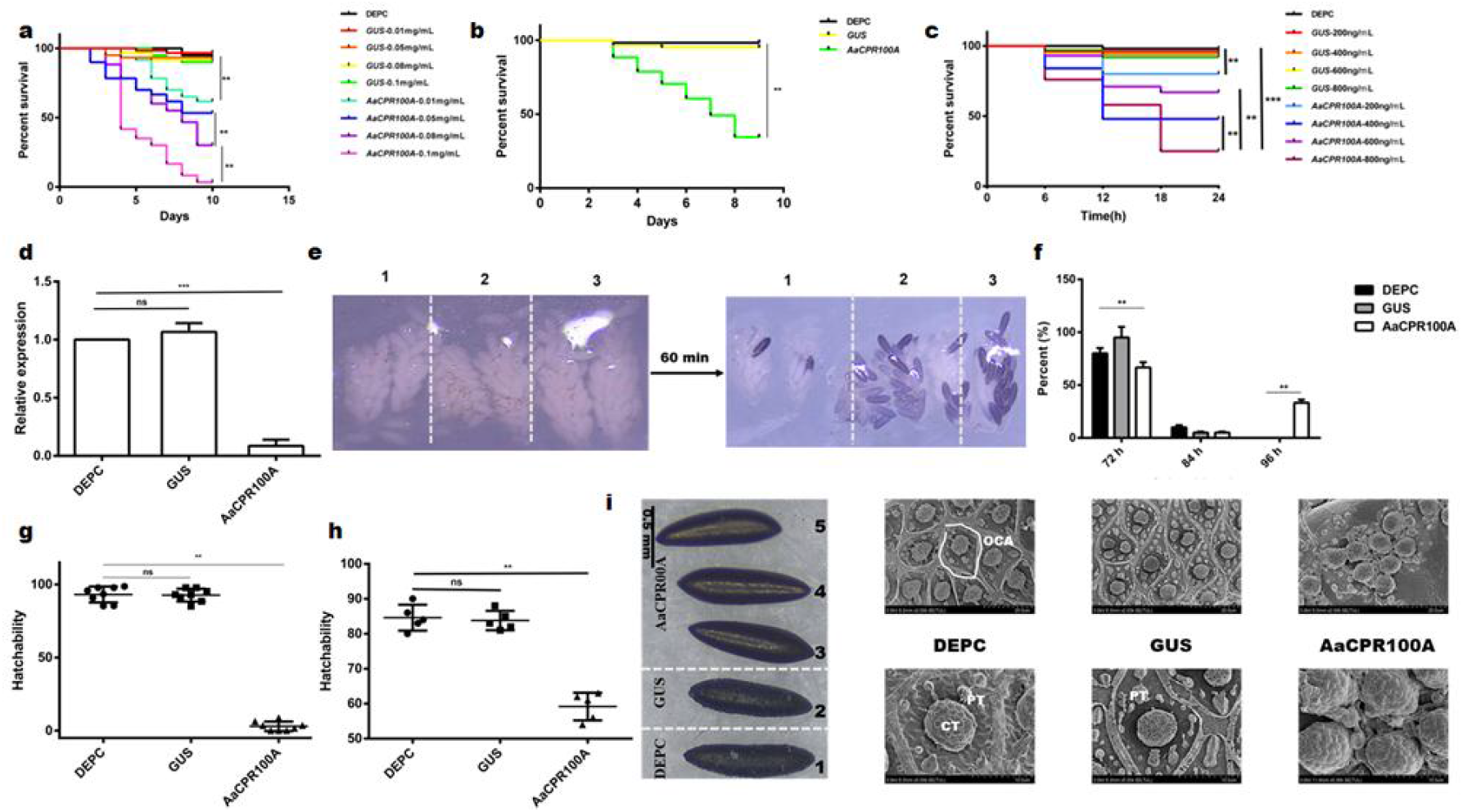
RNAi with *AaCPR100A* in larvae and adults. (a) Survival rate of larvae after immersion with *AaCPR100A* dsRNA. (b) Survival rate of larvae after treatment with bacteria expressing *AaCPR100A* dsRNA. (c) Survival rate of adults after injection of *AaCPR100A* dsRNA. (d) AaCPR100A transcriptional levels at 12 h after injection. (e) Melanization of egg shells at 72 h after blood meal; 60 min is the time after dissection; 1, 2, and 3 in the top of each image represent groups treated with (1) *AaCPR100A* dsRNA, (2) ds*gus* RNA, and (3) DEPC water, respectively. (f) Egg production at different times after blood meal (72 h, 84 h and 96 h after blood meal). (g) Hatchability of eggs laid later. (h) Hatchability of eggs laid at normal time. (i) Optical microscope and SEM images of egg ultrastructure; scale bars are shown in each image. The image on the far left was taken with an optical microscope: 1 is the treatment with DEPC water; 2 is the treatment with ds*gus* RNA; 3–5 are the treatment with ds*AaCPR100A* RNA. The images on the right were taken by SEM: images in the upper panels are in low magnification and images in the bottom panel are in high magnification. Abbreviations: ns, *P* ≥0.05; *, *P* <0.05; **, *P* <0.01; ***, *P* <0.001; AaCPR100A, treatment of larvae and adults with *AaCPR100A* dsRNAs; GUS, treatment of larvae and adults with ds*gus* RNAs; DEPC, treatment of larvae and adults with DEPC water; 0.01 mg/mL and other quantities represent the concentrations of dsRNA. CT central tubercle; PT peripheral tubercles.

Similarly, when AaCPR100A was knocked down, the adult survival rate decreased significantly. Figure 2c shows that when the dsRNA concentration reached 800 ng/mL, only 30% of adults survived to 24 h after injection. RNAi efficiency of the microinjection was also concentration dependent. The surviving adults in the group injected with the highest concentration of dsRNA were selected for qPCR testing. Figure 2d shows that the transcription level of AaCPR100A in the experimental group significantly decreased to 15% (P <0.001). The ovaries were dissected to observe the degree of egg melanization at 48 h after blood meal. The results showed that 70% of eggs in the control groups were melanized at 60 min after dissection, while few of the experimental group were melanized (Figure 2e), which suggests that knockdown of AaCPR100A affected melanization of the chorion.

When AaCPR100A was silenced, the oviposition time of 33.3% of the adults was delayed by 24 h compared with the control groups (Figure 2f), and the hatching rate of eggs produced by the AaCPR100A-silenced mosquitoes was significantly decreased (Figure 2f). Although 55% of the AaCPR100A dsRNA treated mosquitoes laid eggs normally, the egg hatchability decreased significantly compared to the control groups (P <0.0001) (Figure 2g). Figure 2h shows that when the hatching rate of eggs in the control groups was 80%–90%, the hatching rate of AaCPR100A gene knocked down eggs was only 50%. There was no significant difference between treated groups and control groups in the weight of blood meal and the size of eggs (Figure S1). The optical microscope showed that while eggs in the control groups had rough surfaces, AaCPR100A knocked down eggs had relatively smooth surfaces (Figure 2i), which suggests that AaCPR100A participated in the formation of the outer chorionic area (OCA) of eggs, an area that is surrounded by the porous external chorionic network. SEM showed that AaCPR100A affected the formation of the outer chorionic membrane. The central tubercles (CT) converged with each other, and the peripheral tubercles (PT) and outer chorionic network disappeared in the gene knocked down eggs compared to the eggs in control groups (Figure 2i).

### RNAi for adults and TEM for larvae

RNAi treatments using dsRNA on AaCPR100A were fatal in larvae, but the cause of death was not clear. Therefore, adults treated by microinjection of dsRNA were used to observe phenotypic changes in larvae that were hatched from the eggs laid by dsRNA treated adults. Unlike the results of direct RNAi treatment of larvae, the larvae produced by RNAi-treated adults did not show high lethality. In addition, more than 30% of the larvae in the experimental group had lighter pigmentation than those in the control groups (Figure 3a). The RGB (red, green, blue) data of the abdomen and respiratory siphon were extracted by the color picker in Photoshop, and then converted to HSV (hue, saturation, value) values. Darker colors produced smaller vertical values. Figure 3b shows that the larvae from AaCPR100A-dsRNA treated adults had higher values in the abdomen and respiratory siphon, which indicates that the larvae became lighter. TEM showed that the integuments of larvae, whether untreated or treated with dsgus, were normal (Figure 3f, 3g). However, the cuticle microstructure of mosquitoes treated with AaCPR100A-dsRNA had no recognizable endocuticle compared with the mosquitoes in dsgus RNA-treated or untreated groups (Figure 3e).

**Figure 3.**
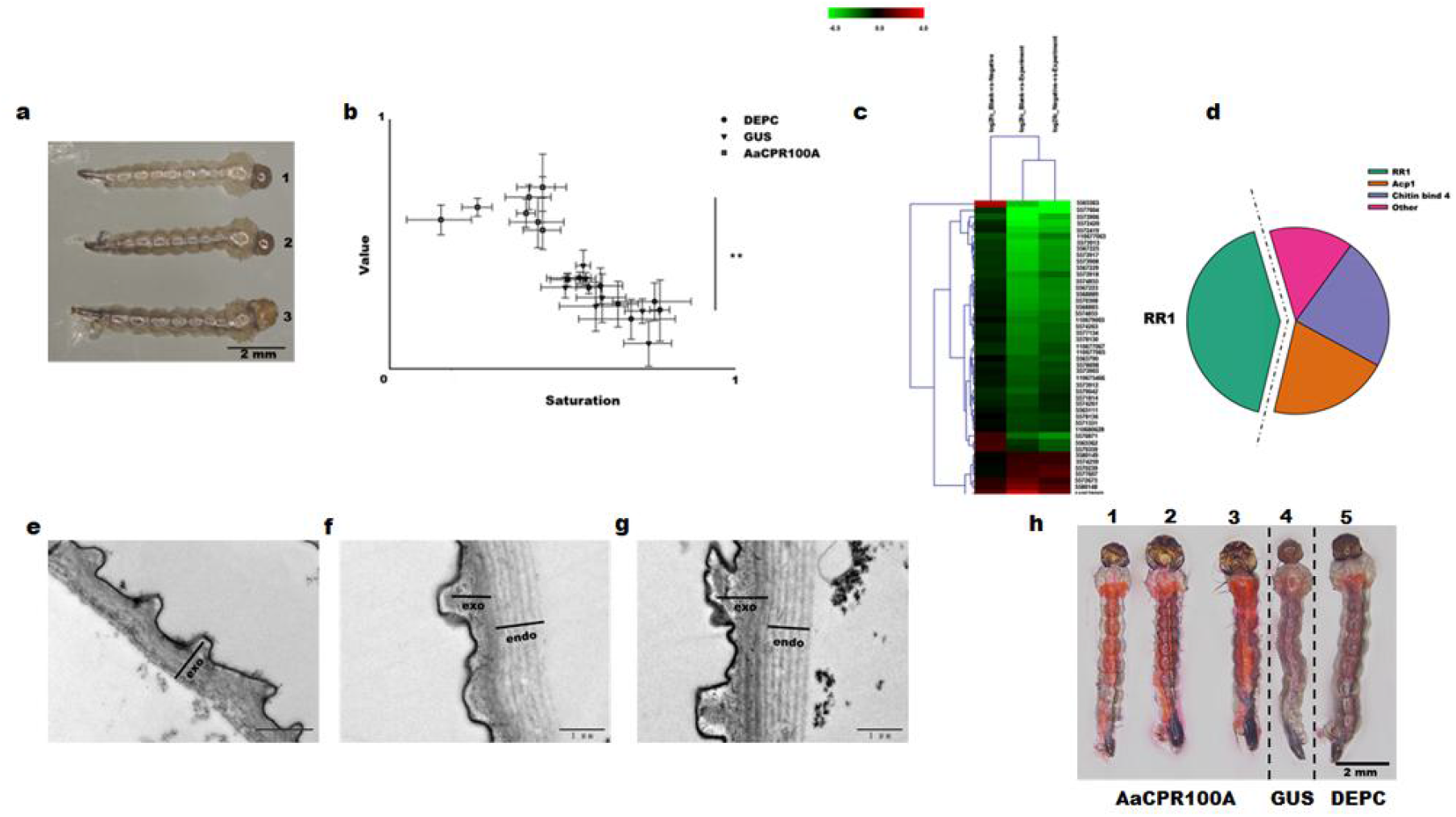
The effect of RNAi of *AaCPR100A* on cuticle formation of larvae. (a) Different level of pigmentation of larvae was observed under an optical microscope; 1, 2 and 3 are the treatments with (1) *AaCPR100A* dsRNAs, (2) *gus* dsRNAs and (3) DEPC water. (b) Differences in larval pigmentation between AaCPR100A dsRNA treatment and control groups for comparison. (c) Cuticular genes that were differently expressed between the *AaCPR100A* knocked down larvae and control group larvae shown by RNA-seq; greater green intensity indicates higher downregulation fold and greater red intensity indicates higher upregulation fold. (d) Types of differential cuticular genes. (e) Integument of larvae treated with AaCPR100A dsRNAs. (f) Integument of larvae treated with *gus* dsRNAs. (g) Integument of larvae treated with DEPC water. (h) Larvae of different treatment groups were soaked with Eosin Y dye: 1–3 are larvae (lighter pigmentation) in the AaCPR100A-dsRNA treatment group; 4 are larvae treated with *gus* dsRNA; 5 are larvae of the blank control. Abbreviations: **, *P* <0.01; AaCPR100A, treatment of larvae with *AaCPR100A* dsRNAs; GUS, treatment of larvae with *gus* dsRNAs; DEPC, treatment of larvae with DEPC water.

### RNA-seq analysis of larvae

RNA-seq analysis showed that 48 cuticle genes were significantly differently expressed compared to the control groups; these 48 genes were not differently expressed between the two control groups (Figure 3c). Genes of the CPR family accounted for 42% of these 48 cuticle genes, and all of them expressed RR1 proteins (Figure 3d). The 33% of larvae with light pigmentation were soaked with Eosin Y dye, and all larvae of the AaCPR100A-dsRNA treated groups that had lighter pigmentation contained more of the Eosin Y dye (Figure 3h).

### Screening and validation of interacting proteins

The average insert size of the cDNA library was > 1200 bp, the library storage capacity was >1.2×107 CFU, the positive rate was 100%, and the library indicators were qualified (Figure S3). AaCPR100A-pGBKT7 were baits. On the four defect plates, only the positive control colony grew and turned blue in color, while the negative control and self-activation colonies did not grow, which indicates that the bait did not self-activate (Figure 4a). After yeast two-hybrid assay, more than 30 proteins which interacted with AaCPR100A were screened from the library, and G12-like was selected for further examination of interaction and function.

**Figure 4.**
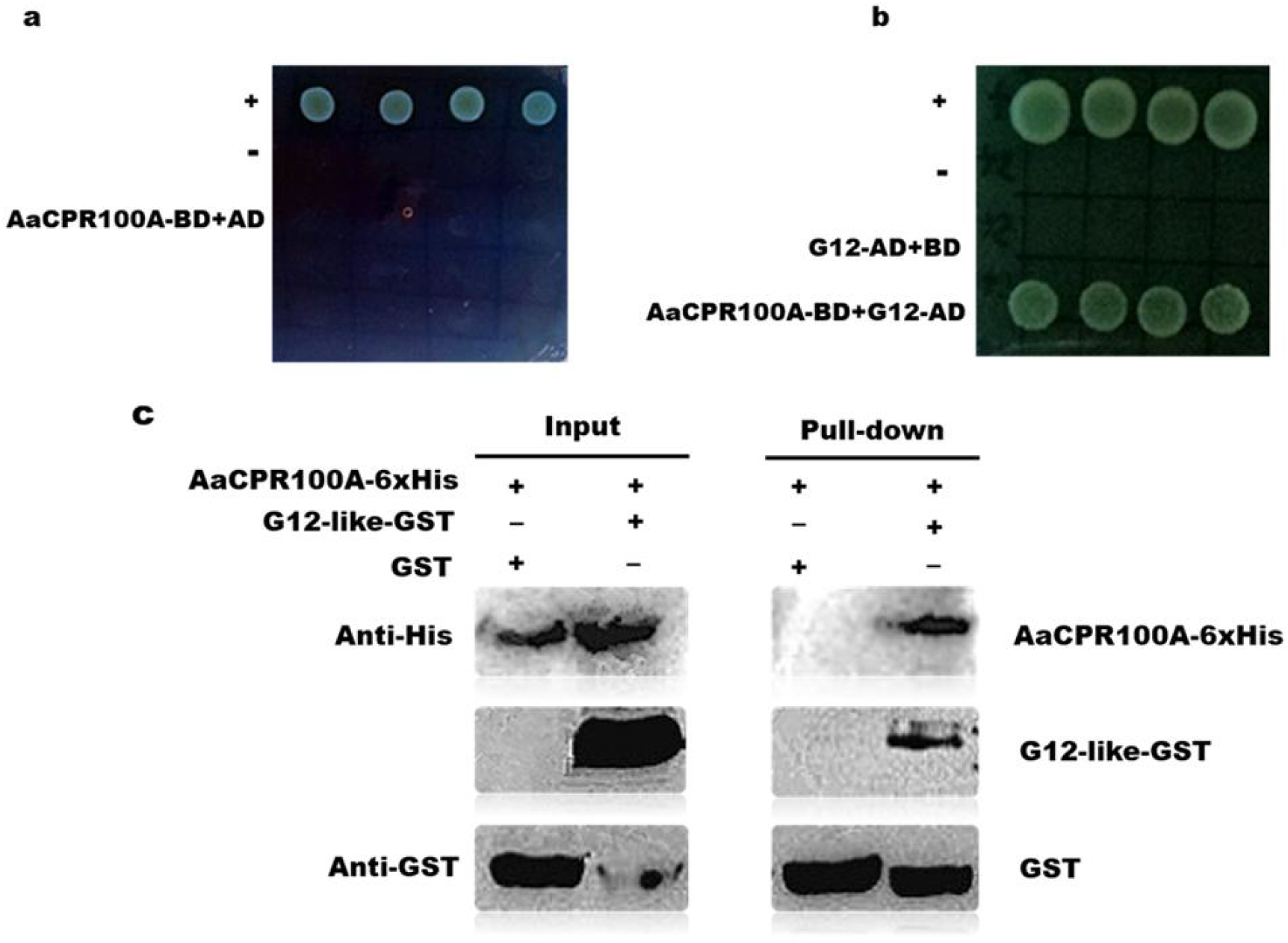
AaCPR100A interaction with G12-like protein. (a) Detection of self-activation of AaCPR100A. (b) Yeast two-hybrid assays showed that AaCPR100A interacts with G12-like. (c) Physical interaction of AaCPR100A and G12-like in vitro was verified by GST pull down assay; GST-G12-like was incubated in binding buffer containing glutathione-agarose beads with or without AaCPR100A-6×His, and the agarose beads were washed six times and eluted. Lysis of *E. coli* (Input) and eluted proteins (pull-down) from beads was immunoblotted using anti-HIS and anti-GST antibodies. Abbreviations: BD, the vector of pGBKT7; AD, the vector of pGADT7.

A recombinant G12-like-pGADT7 sequence was created and transferred together with AaCPR100A-pGBKT7 recombinant plasmid into competent yeast cells that were grown on a SD/-Leu-Trp plate. The G12-like-pGADT7+pGBKT7 vector did not grow and turned green on the SD/-Ade-His-Leu-Trp plate with X-α-gal, while the positive control and the G12-pGADT7+pGBKT7-AaCPR100A vector grew and became green on the four defect plates (Figure 4b), which showed that G12-like protein did not self-activate but did interact with AaCPR100A protein. In vitro physical interaction was observed in GST pull-down assays. AaCPR100A-6×HIS protein was (co-)incubated with and without G12-like-GST protein. Western blot analysis showed that AaCPR100A protein, GST tag and G12-like protein were expressed. Co-incubated proteins and binding buffer containing glutathione-agarose beads were incubated overnight, and the agarose beads were then washed six times and eluted. The pull-down results showed that AaCPR100A protein that was not co-incubated with G12-like protein was not detected in the eluent; only GST was detected. However, in the eluate of AaCPR100A co-incubated with G12-like protein, both AaCPR100A protein and G12-like protein were detected (Figure 4c).

### The expression of *G12-like* and its function in larval survival and growth

G12-like was expressed almost uniquely in the 2nd–4th instar larvae, and its transcription level peaked in the third instar larvae, which is very different from the transcription profile of AaCPR100A (Figure 5a). The expression levels of G12-like in eggs during egg development and in different tissues in adults were also examined, although the transcription levels were very low. The expression of G12-like was greatest at 96-108 h after oviposition (Figure 5b). However, the expression of G12-like was barely detected in any adult tissues before or after blood meal.

**Figure 5.**
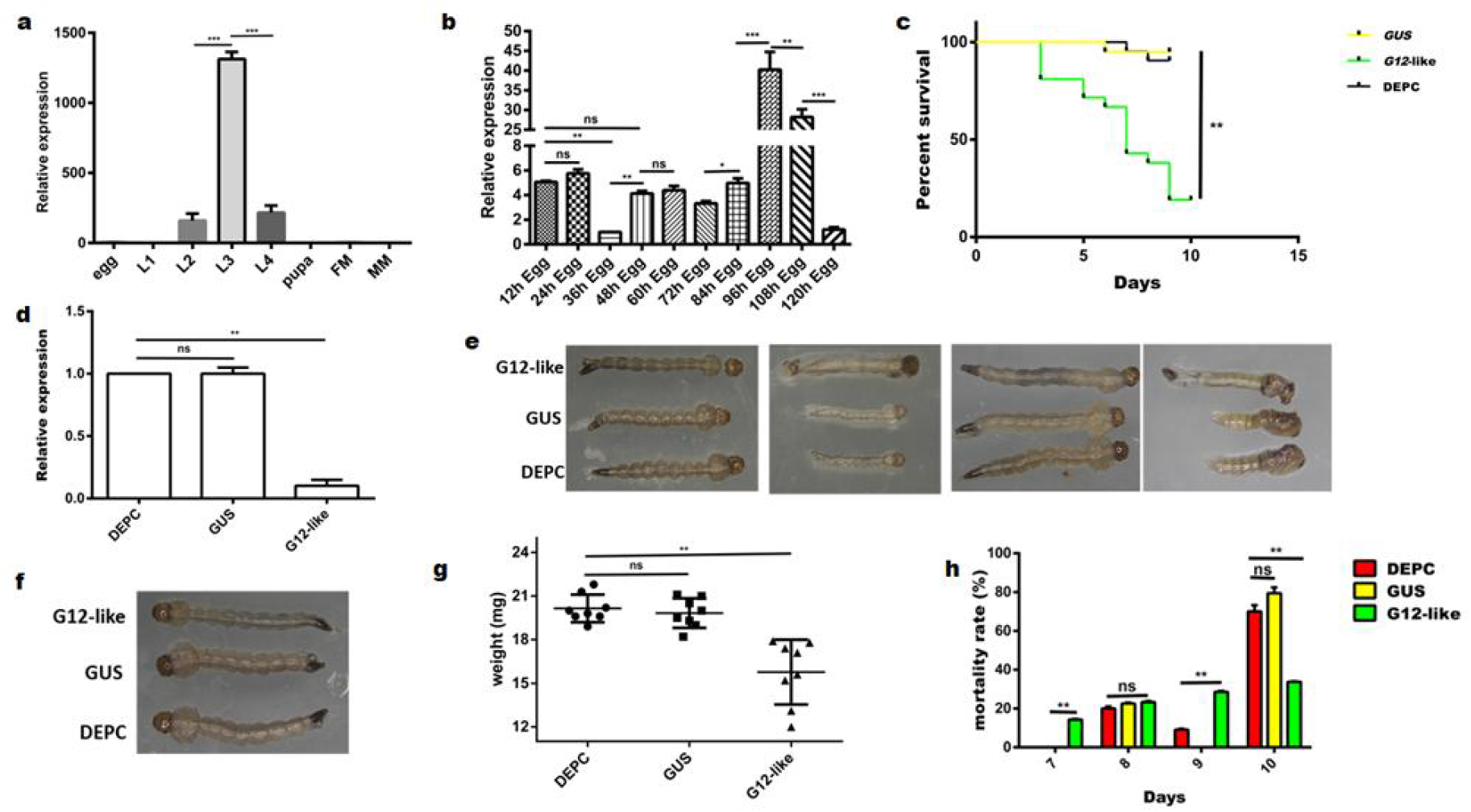
The expression of *G12-like* and its function in larval survival and growth. (a) Temporal transcriptional levels of *G12-like* in different developmental stages. (b) Temporal transcription levels of *G12-like* in eggs with different developmental times; 12 h, 24 h and other times are the times after oviposition. (c) The survival rate of larvae after immersion with *G12-like* dsRNA. (d) *G12-like* transcription levels. (e) The dead larvae fed on *G12-like* dsRNAs were unable to detach from the exuviae. (f) Fourth instar larvae photographed by an optical microscope. (g) The weight of the fourth instar larvae; groups of 5 larvae were weighed together. (h) Low temperature tolerance of larvae; the fourth instar larvae were placed at 4 ºC, and dead mosquitoes were observed every day. Abbreviations: ns, *P* ≥0.05; *, *P* <0.05; **, *P* <0.01; ***, *P* <0.001; L1, first instar larvae; L2, second instar larvae; L3, third instar larvae; L4, fourth instar larvae; FM, female adults; MM, male adults; G12-like, treatment of larvae with dsG12-like dsRNAs; GUS, treatment of larvae with *gus* dsRNAs; DEPC, treatment of larvae with DEPC water.

When the expression of G12-like was knocked down by 75% (Figure 5d), larvae survival rate was significantly low, only 20%, compared to the control groups (Figure 5c). Of the 80% dead larvae, 75% were abnormal in structure, showing molting defects (Figure 5e). The surviving larvae had approximately the same length as the larvae in the control groups (Figure 5f). Figure 5g shows that when the G12-like transcription levels decreased, the weight of the larvae decreased significantly (P <0.001). At the same time, after G12-like knockdown, the larvae in the control groups began to die at 8 d, and 70% of them had died at 10 d; there were no significant differences between the control groups (Figure 5h). However, larvae in the G12-like dsRNAs treated group began to die at 7 d, 24 h earlier than larvae in the control groups (Figure 5h). In addition, mortality of larvae in the G12-like dsRNAs treated group at 8 d and 9 d was greater than that in the control groups, and the remaining larvae died at 10 d (Figure 5h).

### The effect of microinjection of G12-like dsRNAs on adults and eggs

The relative expression of G12-like in the G12-like dsRNAs treated group significantly decreased by 70% (Figure 6a) compared to the control groups. Unlike AaCPR100A RNAi, G12-like expression did not affect the adult survival rate. However, similar to the interference of the gene in larvae, the reduced transcription level of G12-like in adults also led to low temperature intolerance. Figure 6b shows that when adults injected with G12-like dsRNAs were frozen at −20 °C, the mosquito mortality rate was as high as 35%, while almost no mortality was found in mosquitoes in the control groups after the same cold stress treatment.

**Figure 6.**
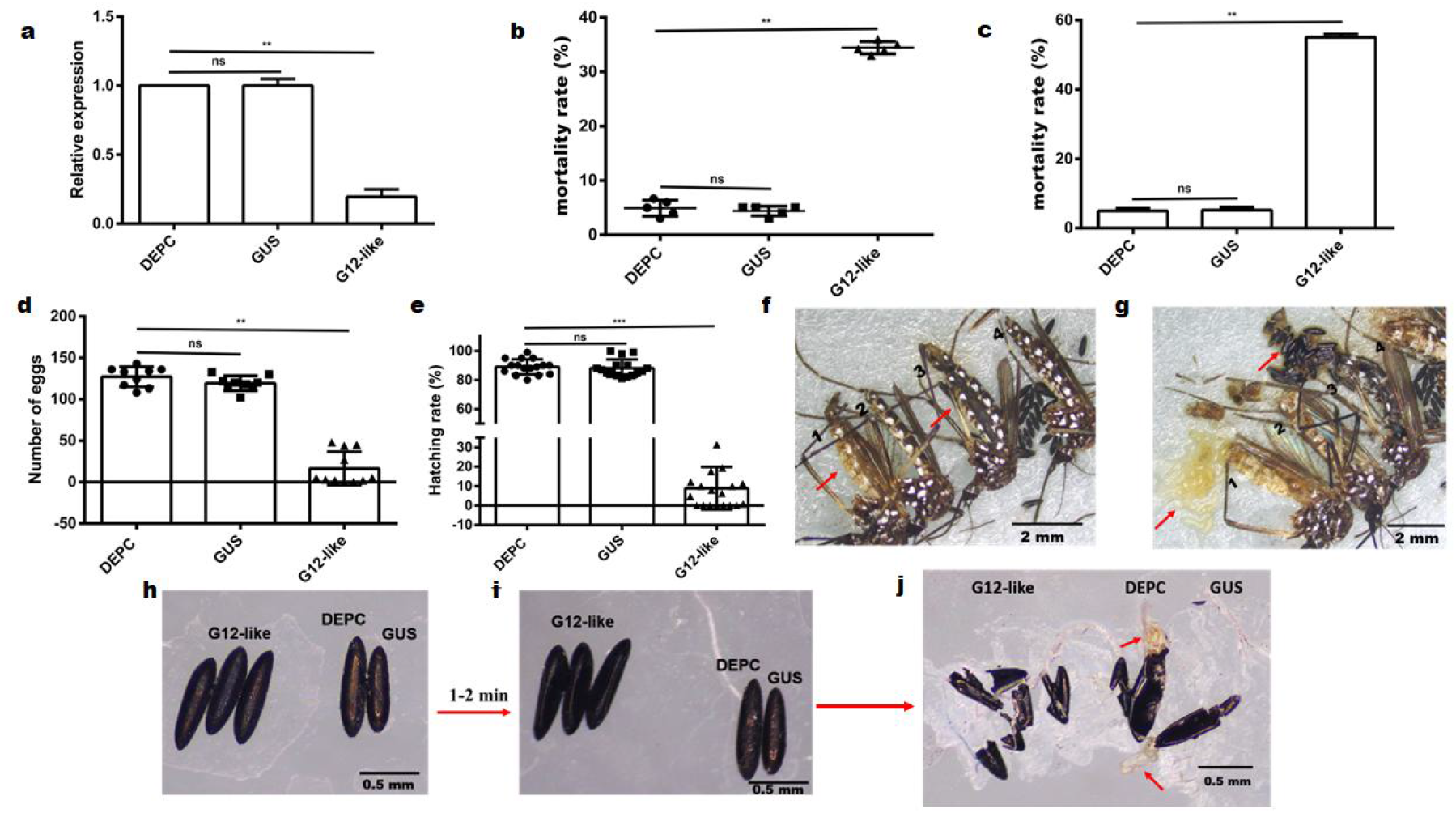
The effect of microinjection of *G12-like* dsRNAs on adults and eggs. (a) *G12-like* transcription levels after different injection treatments. (b) Low temperature tolerance of the adults; tolerance is represented by the mortality rate; higher mortality rate corresponds with less tolerance. (c) Mortality rates of adults in different groups within 3 d after spawning. (d) The number of eggs laid by adults after different injection treatments. (e) Hatching rates of eggs laid in 72 h after different injection treatments. (f) The entire pictures of dystocia adults: 1, an adult mosquito treated with *G12-like* dsRNAs; 2, an adult mosquito treated with *gus* dsRNAs; 3, another adult mosquito treated with *G12-like* dsRNAs; 4, an adult mosquito treated with DEPC water. (g) Dystocia adult mosquitos when the ovary was dissected out: 1, an adult mosquito treated with *G12-like* dsRNAs; 2, an adult mosquito treated with *gus* dsRNAs; 3, another adult mosquito treated with *G12-like* dsRNAs; 4, an adult mosquito treated with DEPC water. (h) Eggs in a humid environment. (i) Eggs in a dry environment. (j) Anatomical location of eggs. Abbreviations: ns, *P* ≥0.05; *, *P* <0.05; **, *P* <0.01; ***, *P* <0.001; G12-like, treatment of adults with *G12-like* dsRNAs; GUS, treatment of adults with *gus* dsRNAs; DEPC, treatment of adults with DEPC water.

The number of eggs laid by G12-like dsRNAs treated mosquitoes significantly decreased (Figure 6d). The number of hatched eggs for each adult mosquito was also counted, which showed that the average hatching rate was 8% for the G12-like dsRNAs treated mosquitoes and the average hatching rate for the control groups was 90% (Figure 6e). There was no significant difference in the weight of blood meal and the size of eggs between G12-like knocked down and control mosquitoes (Figure S4).

Moreover, 54% of G12-like knocked down adults died within 3 d after laying eggs (Figure 6c). The abdomens of adults that had not oviposited were significantly swollen compared with adults in the control groups, and eggs were clearly visible in the abdomens (Figure 6f). It was found that the eggs of the nonovipositing adults had turned yellow or even black in color when dissected out (Figure 6g). Eggs from different treatment groups in the humid environment (36% ±0.2%) were all normal (Figure 6h), but in a dry environment (9.47% ±0.38%), the eggs of the G12-like knocked down group rapidly became desiccated (Figure 6i), while eggs from other groups looked normal. When the eggs were dissected out and carefully checked, we found that there were no developed larvae in the eggs of the G12-like dsRNAs treatment group, while first instar larvae developed from the eggs in other groups (Figure 6j). Maintaining or increasing humidity did not alter this situation. The egg shell surface structure is shown in Figure S5; compared with the control groups, the central tubercles (CT) on the eggshell surfaces in the G12-like interference group were prominent, and the outer chorionic areas (OCA) were blurred.

### Effects of *AaCPR100A* and *G12-like* knockdown on each other’s transcriptional level

The transcription level of each gene in the two genes knocked down mosquitoes was examined to gain further understanding of the interaction between AaCPR100A and G12-like in regulation. The transcription level of G12-like was not affected when the transcription level of AaCPR100A decreased (Figure S6a), but the relative expression of AaCPR100A was downregulated by 45%–50% when the G12-like gene was knocked down (Figure S6b).

### Knockdown of *G12-like* could complement the phenotype and transcriptional abundance of *AaCPR100A* knocked down adults

When AaCPR100A was knocked down, the adult survival rate significantly decreased (Figure 7a); however, knockdown of G12-like did not affect the adult survival rate. When AaCPR100A and G12-like RNAs were interfered simultaneously, the survival rate of adults increased significantly by 80% compared to the RNAi of AaCPR100A alone (Figure 7a). RNAi of AaCPR100A had no significant effect on the transcription level of G12-like (Figure 7b). At the same time, the transcription level of G12-like decreased significantly whether RNAi was of G12-like alone or of G12-like with AaCPR100A simultaneously (Figure 7b). When the expression of G12-like decreased with RNAi, the relative expression of AaCPR100A increased significantly. The expression of AaCPR100A also increased after RNAi of AaCPR100A and G12-like together (Figure 7c).

**Figure 7.**
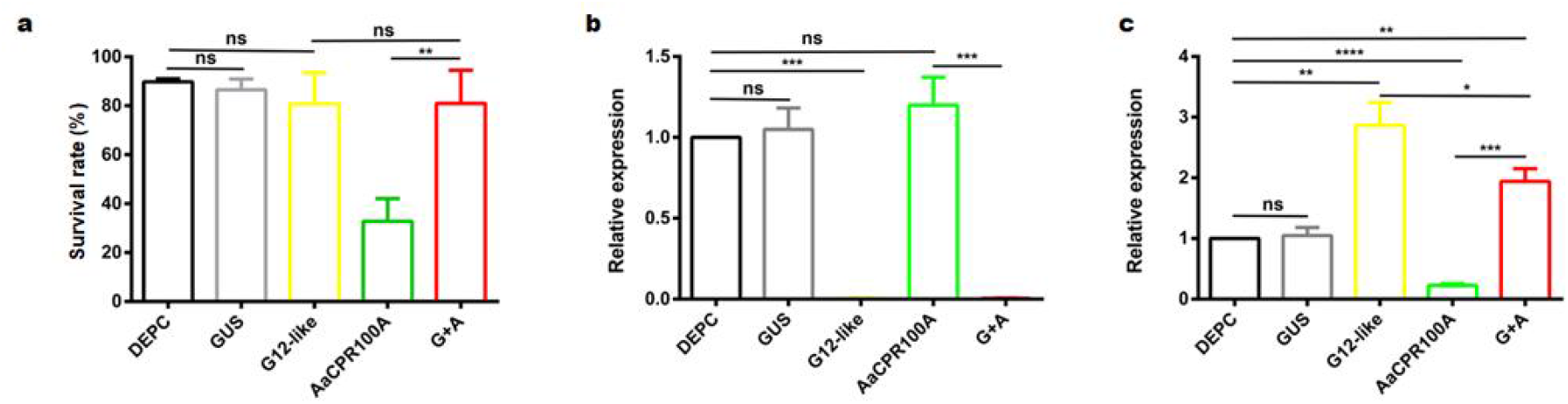
The functional interaction between *G12-like* and *AaCPR100A* was verified by RNA interference. (a) Mortality of adults after RNAi. (b) Changes in the relative expression of *G12-like* after different treatments. (c) Changes in the relative expression of *AaCPR100A* after different treatments. *Abbreviations*: DEPC, treatment of adults with DEPC water; GUS, treatment of adults with *gus* dsRNAs; G12-like, treatment of adults with *G12-like* dsRNAs; AaCPR100A, treatment of adults with *AaCPR100A* dsRNAs; G+A: simultaneous treatment of adults with *AaCPR100A* and *G12-like* dsRNAs; ns, *P* ≥0.05, *, *P* <0.05, **, *P* <0.01, ***, *P* <0.001, ****, *P* <0.0001.

## Discussion

### Lethality of *AaCPR100A* to *Ae. aegypti* by affecting cuticle formation and structure

The cuticle of an insect can be divided from the outside to the inside into envelope, epicuticle, exocuticle, endocuticle and epidermis. The exocuticle and epicuticle together form the procuticle. Lipids exist mainly in the epicuticle. The principal chemical components of the procuticle are chitin and protein, which are secreted during cuticle formation and interact with each other to finally form the orderly arranged cuticle structure *(Moussian, 2010)*. AaCPR100A belongs to the CPR gene family and contains the RR1 motif, which can be preliminarily classified as a flexible cuticle protein. Our immunohistochemical analysis showed that the AaCPR100A protein may be abundant in the female adult mosquito thorax, fat body and intersegment membranes. The qPCR analysis indicated that the transcription level of AaCPR100A was greatest in eggs, followed by the thorax and fat body. These results suggest that AaCPR100A expression is crucial in the development of the soft organs and tissues of the adult body as well as in regions with cuticle protection.

Both larvae and adults showed strong evidence of lethality when AaCPR100A was knocked down, but the proximate cause of death was not clear. Larvae also clearly showed abnormal pigmentation on the body surface and the absence of endocuticle. The RNA-seq data of these larvae showed that when AaCPR100A expression decreased, transcription of some cuticular genes was significantly affected; 50% of these genes were in the CPR family, and all of them transcribed the RR-1 motif. Vannini and Willis (15) found that that two forms of the consensus, RR-1 and RR-2, have been recognized. Initial data in their study suggested that RR-1 and RR-2 proteins were present in different regions within the cuticle, and that RR-2 proteins contributed to the formation of the exocuticle, which becomes sclerotized, while RR-1s was found in the endocuticle, which remains soft *(Vannini and Willis, 2017)*. Our results suggest that AaCPR100A is important in the formation of endocuticle and flexible cuticles, and the direct interference of AaCPR100A in larvae resulted in high lethality. This lethality may have been due to the influence of the development of the cuticle barrier during exfoliation or the hardening, blackening and permeability of the cuticle after molting.

### The upstream interaction of protein G12-like with AaCPR100A promotes its action in larvae

We identified that AaCPR100A interacted with G12-like protein, and RNAi with them in larvae resulted in high lethality. RNA-seq results also showed that the transcriptions of some cuticle, chitin and lipid metabolism-related genes were affected (Figure S2). Larval growth and cold temperature resistance also significantly decreased in G12-like knocked down larvae. We hypothesize that interaction between AaCPR100A and G12-like proteins may also affect fat synthesis on Ae. aegypti. Our findings here suggest that the upstream interacting protein G12-like regulate AaCPR100A because when AaCPR100A was knocked down in larvae, transcription of G12-like was not affected, but when G12-like was knocked down, transcription of AaCPR100A was downregulated.

We also investigated the function of G12-like on mosquitoes. RNAi results showed that when the G12-like expression level was knocked down, the mortality was lower. And there was a difference from larvae treated with AaCPR100A dsRNAs: the cause of death in G12-like knocked down larvae was relatively clear, with most dying from molting abnormalities. A previous study showed that in the final larval stage, the larva must be above a minimal viable weight for metamorphosis. The minimal viable weight was defined as the minimal weight at which the amount of fat body storage is sufficient for survival through metamorphosis *(Mirth and Riddiford, 2007)*. In the last larval stage of Manduca sexta, failure to exceed minimum viable weight impaired metamorphosis *(Nijhout, 1975)*. In addition, the decreased food intake in dsInR-injected larvae prevented the larvae from reaching the minimum viable weight; i.e., they had insufficient nutrient storage in their fat body to survive through metamorphosis *(Lin et al., 2016)*. In the study, the weight of larvae treated with G12-like dsRNAs significantly decreased, and the larvae showed intolerance of cold temperature, which suggested that abnormal exfoliation death was associated with the influence of fat metabolism required for providing molting energy.

### The upstream interacting protein G12-like of AaCPR100A inhibits its function in adults

Knockdown of AaCPR100A caused great mortality, but knockdown of G12-like was not fatal in adult mosquitoes. However, the mosquitoes that survived AaCPR100A knockdown showed no obvious intolerance to cold temperature or any other abnormal phenotypes, while G12-like knocked down mosquitoes showed obvious intolerance to cold temperature. These findings suggest that some deaths of G12-like knocked down mosquitoes within 3 d after egg deposition may be associated to lack of energy caused by abnormal synthesis of fatty acid. We found, by carefully reviewing the functions of any homologs of the G12-like gene in other organisms, G12-like is possibly involved in fatty acid biosynthesis. It has been found that midline-1 interacting with a G12-like protein, MIG12, was recently identified as an acetyl-coenzyme A carboxylase-binding protein in mammals *(Inoue et al., 2011)*. The binding induces de novo fatty acid (FA) synthesis through the activation of acetyl-coenzyme A carboxylase (a rate-limiting enzyme for de novo fatty acid synthesis) (Berti et al., 2004; Inoue et al., 2011).

When the transcript abundance of G12-like decreased, the relative expression of AaCPR100A was upregulated in adult mosquitoes. However, when AaCPR100A was knocked down by RNAi, the transcription abundance of G12-like was not affected. And the expression of AaCPR100A was upregulated in adult mosquitoes in which both AaCPR100A and G12-like knocked down by RNAi. When only AaCPR100A was knocked down, the survival rate of adults was 35%, but when both AaCPR100A and G12-like were knocked down, the survival rate of adults increased to 80%. These results suggest that G12-like both inhibits the transcription abundance of AaCPR100A in adults and decreases the death rate of adults caused by AaCPR100A knockdown. In all, the results demonstrate that G12-like inhibits AaCPR100A and influences its upstream transcription. However, the underlying mechanism by which G12-like affects AaCPR100A remains unclear and is a future research topic.

### AaCPR100A and its interacting protein G12-like affect eggshell integrity

The hatching rates of eggs laid by AaCPR100A knocked down mosquitoes decreased significantly due to loss of egg shell integrity. The hatching rates of eggs laid by G12-like knocked down mosquitoes also significantly decreased. Furthermore, mature eggs were desiccated and shriveled rapidly in a dry environment. Melanin promotes the formation of the serosal cuticle that protects eggs from desiccation *(Farnesi et al., 2017)*. We found that eggs laid by G12-like knocked down mosquitoes had no developing larvae and shriveled rapidly in a dry environment. Lipids constitute approximately 35% of the dry weight of Ae. aegypti eggs; most of the lipids are derived from fat body and are transported to the ovaries through the hemolymph *(Chung et al., 2014)*. We therefore inferred that when de novo synthesis of FA in the eggs was blocked. FA content was insufficient for the eggs to develop normally, and the consequent abnormal development did not initiate functions necessary to protect eggs from desiccation and caused abnormal ovarian development after maternal blood meal. Thus, some eggs became deformed in the ovary and could not be laid.

## Conclusion

In summary, our results indicate that AaCPR100A is an essential gene in Ae. aegypti. AaCPR100A together with its interaction protein G12-like affects the molting, desiccation resistance, and cold temperature tolerance of Ae. aegypti. G12-like is an upstream interaction protein of AaCPR100A at the transcriptional level, and is involved in cuticle and egg shell formation. It is also stage specific and may act in synergy with AaCPR100A in the larval stage and inhibit AaCPR100A in the adult stage, although we lack a full understanding of its mechanism (Figure 8).

**Figure 8.**
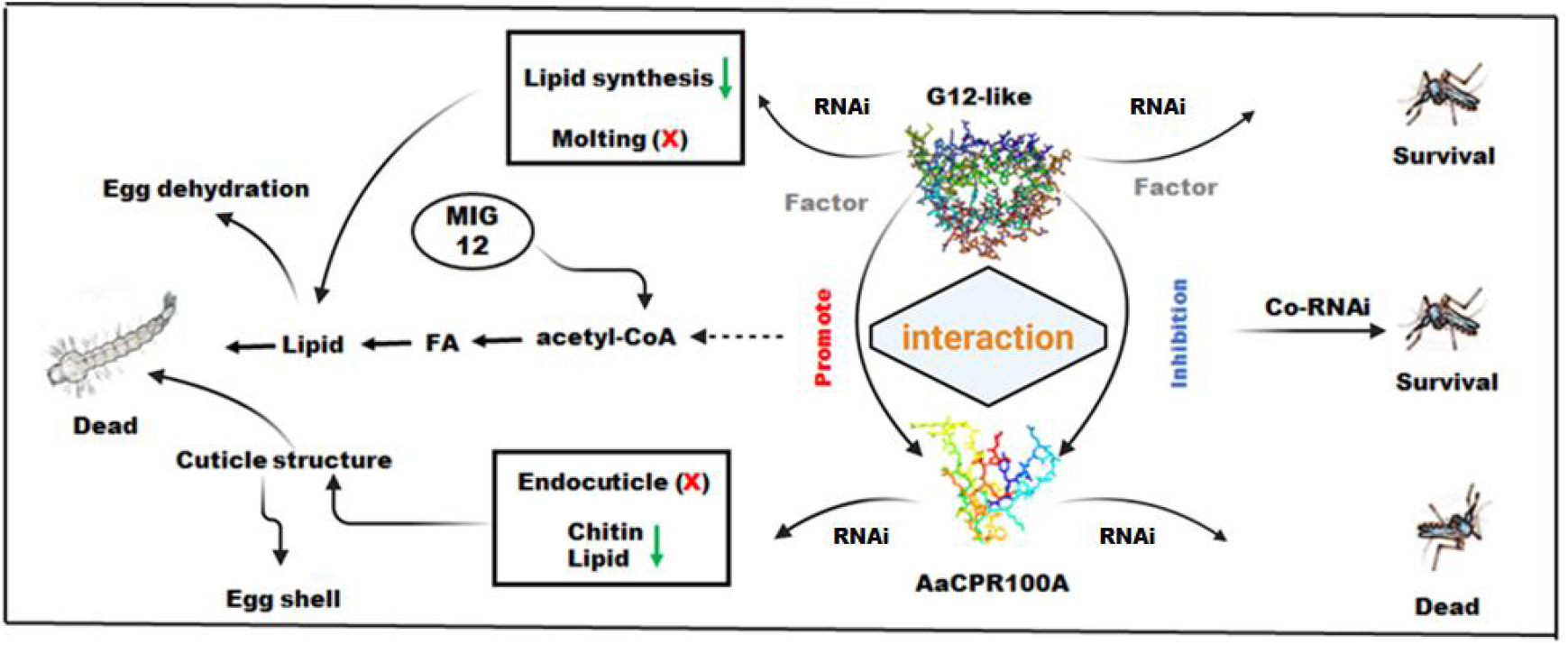
Model of the proposed mechanism of the interaction between lethal cuticular structural protein AaCPR100A and its upstream interacting protein G12-like and their functions. AaCPR100A affects the formation of the endocuticle of larvae and the integrity of the egg shell. The upstream interaction protein G12-like promoted AaCPR100A function in larvae, but inhibit the role of AaCPR100A in adults. We speculate that there may be other factors involved in the regulation of AaCPR100A by G12-like.

## Methods

### Mosquito rearing

*Ae. aegypti* (Rockefeller strain, provided by Beijing Institute of Microbiology and Epidemiology, Beijing, China) was used in the investigation. Mosquitoes were reared according to the previously reported method (44).

### Quantitative Real-Time PCR (qRT-PCR)

Total RNA was extracted using Trizol reagent (Invitrogen, USA), and cDNA was synthesized using PrimeScript™ RT reagent kit with gDNA Eraser (TaKaRa, Dalian, China). Primers used for qPCR (forward: 5′-ACA GTA CGA TCC TCG TTG GA-3′ and reverse: 5′-GGC GGT GAT AGT TTT CGA TT-3′) were designed to amplify fragments of *AaCPR100A* (XM_001662924.2). And the primers (forward: 5′-GTCCAACGAAGCCAATA-3′ and reverse: 5′-GAA ACC ATC CGA ACA GT-3′) were designed to amplify fragments of *G12-Like* (XM_021841051.1). *GAPDH* (glyceraldehyde 3 phosphate dehydrogenase) (XM_011494724) was used as an internal reference gene with primers (forward: 5′-TTG GAC TAC ACC GAA GAG GA-3′ and reverse: 5′-TGT CGT ACC AGG AGA TGA GC-3′). Then the mRNA expression was analyzed quantitatively using the SYBR Green Supermix (Roche, Basel, Switzerland), and the relative gene expression data were analyzed using the relative 2^−ΔΔct^ method (45).

### DsRNA preparation and gene silencing

The primers (forward: 5′-CCG CTC GAG TAG CGG GCT AGT GCC CAG TAT A-3′ and reverse: 5′-AAA AGG AAA ACG CCG GCG AAG TCT GTC GTT GTG AGG GCA-3′) used for *AaCPR100A* RNAi were designed by Primer 5 based on the sequence coding Pfam domain. β-glucuronidase (*gus*) gene (KY848224), a bacterial gene specific to *Escherichia coli*, as a negative control, was amplified by PCR from the pBI121 plasmid using the primers (forward: 5′-GCG GCC GCC CCT TAC GCT GAA GAG ATG C-3′ and reverse: 5′-CTC GAG GGC ACA GCA CAT CAA AGA GA-3′) as reported. And the primers (forward: 5′-ATT TGC GGC CGC TTT AGA AAC TGC TCT CGC TGT-3′ and reverse: 5′-CCG CTC GAG CGG ATC GGT ATC GGT AAC GG-3′) used for *G12-like* were also designed by Primer 5. These three PCR products were cloned into the dsRNA transcription plasmid pL4440, to be used to transform *E. coli* DE3 (HT115) (Weidi, Shandong, China). The experiments were performed according to a previously published method (46).

### Yeast two-hybrid assays

The target gene *AaCPR100A* was constructed into plasmid pGBKT7, the primers (forward: 5′-CGC CAT ATG GCG AGT ATA ATG GAT AAA CCG T-3′ and reverse: 5′-CCG GAA TTC CGG AAC CTT GTT GAA GGT GTA GC-3′) were used to amplify *AaCPR100A*. The constructed plasmid was then transformed with pGADT7 by yeast strain AH109, and verified whether it has self-activation reaction through synthetic dropout (SD)-Ade-His-Leu-Trp (Clontech, CA, USA) with X-α-galactosidase (x-α-gal). Subsequently, the yeast strain AH109 was used to co-transform pGBKT7-*AaCPR100A* bait plasmid and library plasmid simultaneously. Transformed yeast cells were assayed for growth on (SD)/-Trp-Leu plates and SD/-Trp-Leu-HisAde plates containing X-a-gal.

### Validation of G12-like and AaCPR100A protein interaction by yeast two-hybrid assays

According to the sequencing results, the full length of the corresponding library plasmid gene was obtained through a sequence comparison in NCBI. To construct a recombinant plasmid *G12-like*-pGADT7, primers (forward: 5′-CGC CAT ATG GCG ATG AAA CTG CTC T – 3′ and reverse: 5′-CCG GAA TTC CGG CTA GTT CCA CTG GAA GAA TC-3′) were used to amplify *G12-like* gene. The plasmid *G12-like*-pGADT7 and the bait protein gene plasmid *AaCPR100A*-pGBKT7 were co-transformed into competent AH109 yeast cells, coated with SD-Leu/-trp (Clontech, USA) plate, and at the same time, negative and positive controls were also set. The cells were cultured at 30 ℃ for 1-3 days. Some single colonies were selected and placed on SD-Ade-His-Leu-Trp plates containing x-α-gal, and cultured at 30 °C for 1-3 days to observe whether the colonies grew and turned blue.

### In vitro protein-protein interaction assay-glutathione S-transferase (GST) pull-down

pCold TF, *AaCPR100A*-pCold TF, pGEX-6p-1 and *G12-like*-pGEX-6p-1 recombinant plasmids were introduced into the *E. coli* strain BL21 (DE3), and the expression of proteins was induced with 0.6 mM isopropyl b-D-1-thiogalactopyranoside (IPTG) at 16°C for 22 h. The primers (forward: 5′-CGC CAT ATG GCG AGT ATA ATG GAT AAA CCG T-3′ and reverse: 5′-CCG GAA TTC CGG AAC CTT GTT GAA GGT GTA GC-3′) and the primers (forward: 5′-CGC CAT ATG GCG ATG AAA CTG CTC TCG CTG TT-3′ and reverse: 5′-CCC CTC GAG GGG GTT CCA CTG GAA GAA GAA TCC AC-3′) were used to amplify *AaCPR100A* and *G12-like*, respectively. Equal amounts of GST-*G12-like* and His-*AaCPR100A* sonicated lysates were mixed with high-affinity GST resin (GenScript, Piscataway, NJ, USA), and then eluted. Next, the proteins were separated by 12 % SDS-PAGE and immunoblotted with anti-GST or anti-His antibody (Abcam, Cambs, England).

### Scanning electron microscopy (SEM)

Specimens were fixed in 2.5 % glutaraldehyde (Kemiou, Tianjin, China) overnight, and samples were washed 3-4 times for 15 min with phosphate buffer (0.1 M, pH 7.2). Specimens were further fixed with 1 % osmium tetroxide buffer at 4 °C for 1–2 h. After fixation, specimens were washed 3-4 times with distilled water for 15 min. The samples were gradually dehydrated with 30, 50, 70, 90, 95 and 100 % ethanol for 5 to 10 min at each step; the 100 % ethanol step was repeated 2 to 3 times to ensure complete dehydration. After the samples were dehydrated, added isoamyl acetate, overnight at 4 °C. Platinum film was plated with JFC-1600 (Jeol, Tokyo, Japan). The samples were observed using a field emission scanning electron microscope SU8020 (Hitachi Limited, Tokyo, Japan).

### Transmission electron microscopy (TEM)

Specimens were fixed in 2.5 % glutaraldehyde in phosphate buffer (0.1 M, pH 7.0) overnight, and samples were washed 3 times for 5 min with phosphate buffer. Specimens were further fixed with 1% osmium tetroxide buffer in phosphate buffer (0.1 M, pH 7.0) at 4 °C for 1–3 h. After fixation, specimens were washed 3 times with phosphate buffer (0.1 M, pH 7.0) for 5 min. The samples were gradually dehydrated with 30, 50, 70, 90, 95 and 100 % ethanol for 5 to 10 min at each step; the 100% ethanol step was repeated 2 to 3 times to ensure complete dehydration.

After dehydration, the samples were embedded in epoxy resin (Sigma-Aldrich, St. Louis, USA). The specimens were sliced with an ultramicrotome (Leica UC6; Leica, Vienna, Austria). The sections were stained with uranyl acetate (Syntechem, Changzhou, China) and lead citrate (Acros, Shanghai, China) and observed using an electron microscope (JEM1200EX; Jeol, Tokyo, Japan).

## Acknowledgments

We thank Dr. Michael R. Strand (Department of Entomology, University of Georgia, Athens, GA 30602, USA) for English corrections, suggestions and comments on an earlier draft of the manuscript. This study was supported by the National Natural Science Foundation of China (31960703, U22A20363) and the Major Science and Technology plan of Hainan Province (ZDKJ2021035).

## Author Contributions

CL and QH conceived the general hypothesis; CL and JC designed the experiments; JC, YW, HL carried out the experiments; JC, CL, QH, GC and ZT analyzed the results and wrote the paper.

## Competing Interest Statement

The authors declare no competing financial interests.

## Notes

### Competing Interest Statement

The authors have declared no competing interest.

