## Supplementary figures and images for "A novel lethal cuticular structural protein, AaCPR100A and its upstream interaction protein, G12-like, function in cuticle and egg shell formation in the yellow fever mosquito, *Aedes aegypti*"

### Supplemental Fig. 1

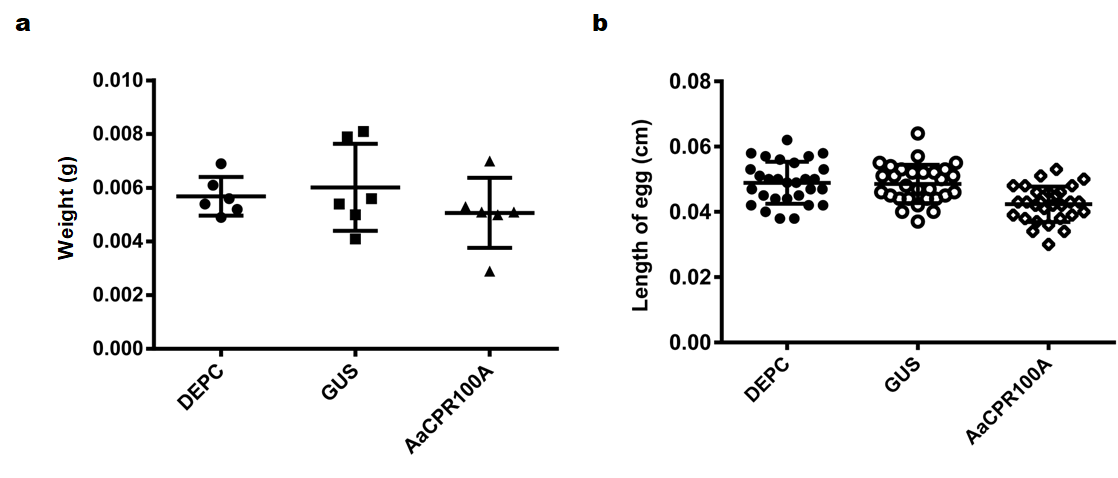

### Supplemental Fig. 2

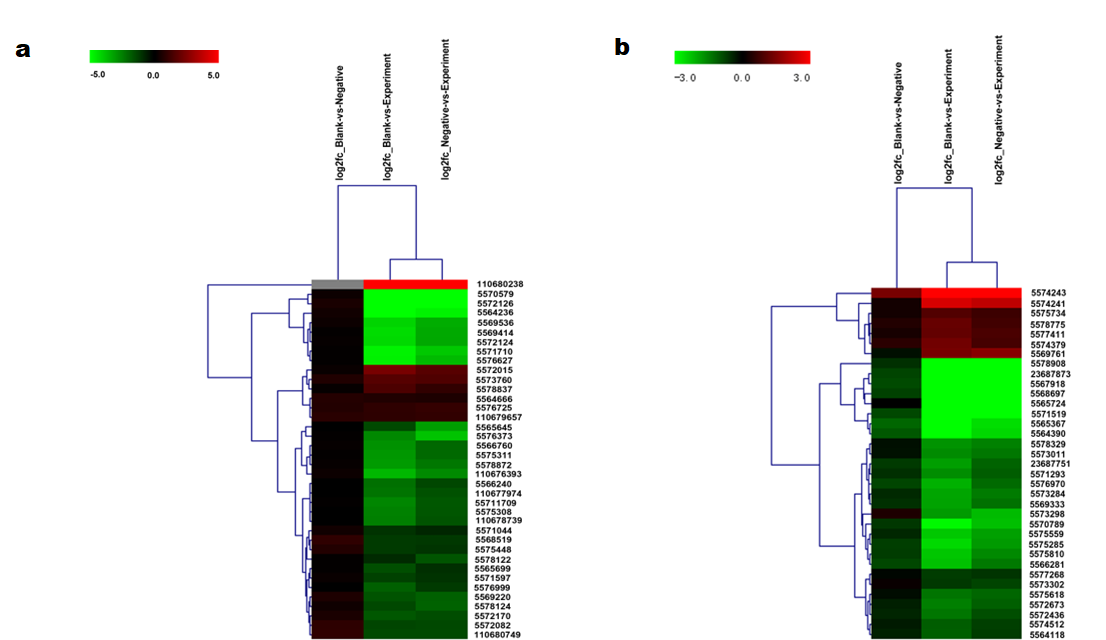

### Supplemental Fig. 3

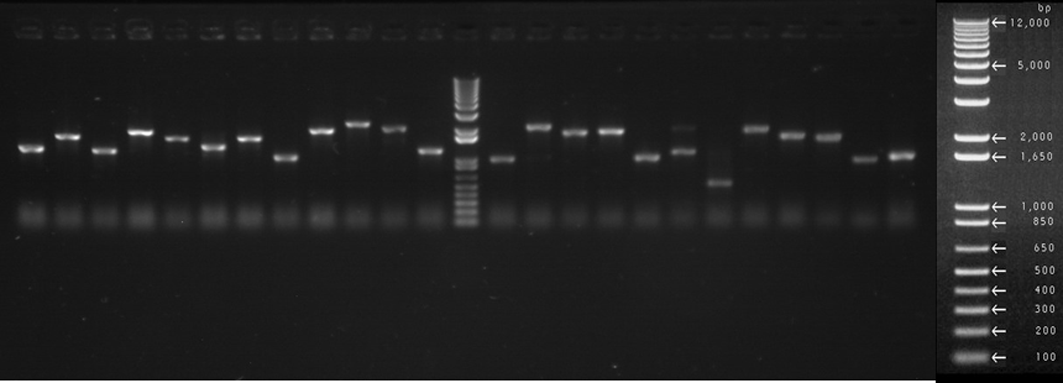

### Supplemental Fig. 4

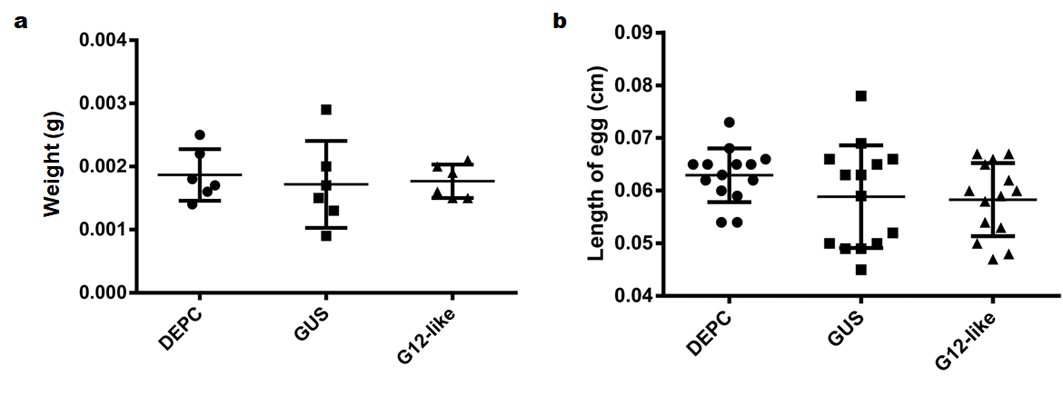

### Supplemental Fig. 5

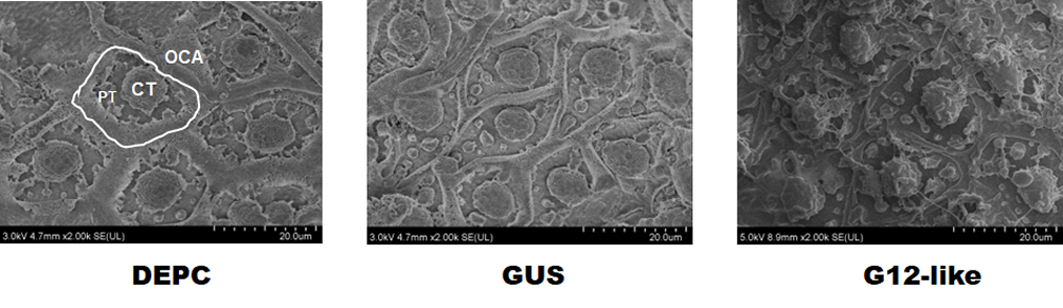

### Supplemental Fig. 6

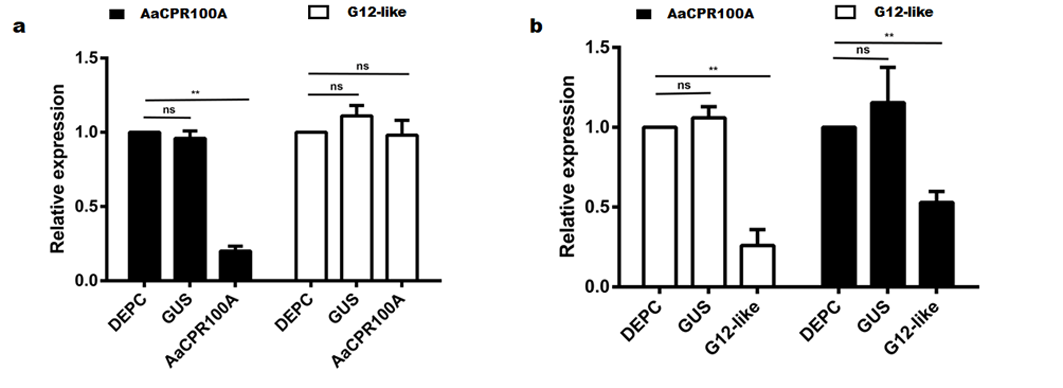
